# Ubiquity of collective irregular dynamics in balanced networks of spiking neurons

**DOI:** 10.1101/389452

**Authors:** Ekkehard Ullner, Antonio Politi, Alessandro Torcini

## Abstract

We revisit the dynamics of a prototypical model of balanced activity in networks of spiking neurons. A detailed investigation of the thermodynamic limit for fixed density of connections (massive coupling) shows that, when inhibition prevails, the asymptotic regime is not asynchronous but rather characterized by a self-sustained irregular, macroscopic (collective) dynamics. So long as the connectivity is massive, this regime is found in many different setups: leaky as well as quadratic integrate-and-fire neurons; large and small coupling strength; weak and strong external currents.

---

Dynamical regimes where excitation and inhibition almost balance each other are considered very important in computational neuroscience, since they are generically accompanied by strong microscopic fluctuations such as those experimentally observed in the resting state of the mammalian brain. While much is known on the balanced regime in the context of binary neurons and in networks of rate models, much less is known in the more realistic case of spiking neural networks. So far, most of the research activity on spiking neurons was restricted to diluted networks, with the goal of providing a detailed description of the underlying asynchronous regime. In this paper we show that, contrary to the current expectations, even in the presence of a 10% dilution, the collective dynamics exhibited is characterized by a sizeable synchronization. The analysis of a suitable order parameter reveals that the macroscopic dynamics is highly irregular and remains such in the thermodynamic limit (i.e. for infinitely many neurons). The underlying form of synchronization is thereby different from the collectively regular dynamics observed in systems such as the Kuramoto model.

In spite of the many studies carried out in the last decade, a general theory of the dynamics of large ensembles of oscillators is still lacking even for relatively simple setups where the single units are assumed to be one-dimensional phase oscillators [1]. A whole variety of phases has been indeed discovered which interpolate between the fully synchronous and the asynchronous regime, including chimera states, self-consistent partial synchrony, not to speak of various clustered states [2–5].

Even though real systems are composed of a finite number of elements, we know from statistical mechanics that a meaningful identification of the different regimes can be made only in the thermodynamic limit, i.e. for an ideally infinite number of elements. In the case of dynamical systems defined on regular lattices with short range interactions, taking the limit is straightforward: it is just the matter of considering infinitely extended lattices. In networks with long-range interactions, the question is less obvious [6]. Since the interaction grows with the system size, the coupling strength must be inversely proportional to the number of connections to avoid unphysical divergencies. Systems like the Kuramoto model belong to this class [2]. In setups where the average coupling contribution is negligible, the coupling strength is instead assumed to scale as the inverse of the square root of the connectivity. Spin glasses are the most prominent physical systems where this latter scheme is adopted [7, 8].

The characterization of the balanced regime represents another such setup [9] encountered in computational neuroscience. A theory of balanced states has been developed in ensembles of neurons characterized by a coarsegrained variable: their firing-rate [10–15]. However, it is still unclear whether the resulting scenario is truly representative of what can be observed in more realistic setups.

In fact, an increasing attention has been progressively devoted to simple models of excitatory and inhibitory spiking neurons, such as leaky (LIF) or quadratic (QIF) integrate-and-fire neurons with the goal of mimicking the cortical activity [16–20]. The most detailed theoretical analysis of spiking neurons has been proposed by Brunel [21], who derived and solved a (self-consistent) Fokker-Planck equation for the probability density of membrane potentials in a network of LIF neurons. The theory was developed by assuming a finite sparse connectivity, so that the thermodynamic limit is implicitly taken by letting the number of neurons diverge. As the resulting scenario - an asynchronous regime and two kinds of synchronous activity - does not fully match the one found in rate models, several numerical studies have been performed to investigate the role of ingredients such as the synaptic time scale or the network connectivity [11–14, 22]. The overall result is the evidence of some features which seem to conflict with the hypothesis of a widespread existence of a single “standard” asynchronous dynamics. For instance Ostojic claims the existence of two different regimes that can be detected upon increasing the coupling strength [23]. Even though this statement has been challenged by Engelken et al. [24], who maintain that a single, standard, asynchronous regime does exist, the qualitative features of the spiking activity need to be better understood.

In this Article, we revisit the activity of a balanced network of spiking neurons and propose a different interpretation of the regimes that have been observed in simulations of finite networks. Our approach is based on a thermodynamic limit, which better preserves the qualitative features observed in finite systems. All of our studies show that the network activity is not asynchronous but rather a manifestation of a collective irregular dynamics (CID), similar to what found in heterogeneous networks of globally coupled inhibitory neurons [25].

Collective dynamics can be quantified by implementing the same indicators introduced to measure the degree of synchronization. With the help of an order parameter specifically designed to characterize neuronal synchrony in large ensembles of neurons [26], we find that CID is an ubiquituous phenomenon, which does not only persist for arbitrary coupling strength, but also in the absence of delay and refractoriness. Finally, we find that collective dynamics is not restricted to LIF neurons, but extends at least to QIF neurons as well. All numerical calculations have been performed by implementing either an event-driven approach [27, 28] or Euler’s algorithm.

We start considering an ensemble of N supra-threshold LIF neurons composed of *bN* excitatory and (1 − *b*)*N* inhibitory cells, as defined in Refs. [21, 23]. The membrane potential *V_i_* of the *i*th neuron evolves according to the equation,

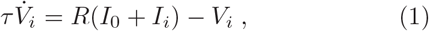

where *τ* = 20ms is the membrane time constant, *RI*_0_ = 24mV is an external DC “current”, and *RI_i_* the synaptic current arising from the mutual coupling

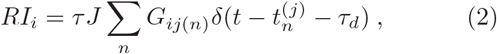

where *J* is the coupling strength. The synaptic connections among the neurons are random, with a constant indegree *K* for each neuron. The matrix elements assume the following values: *G_ij_* = 1 (−*g*), if the pre-synaptic neuron *j* is excitatory (inhibitory), otherwise *G_ij_* = 0. If *V_j_* reaches the threshold *V_th_* = 20 mV at time 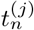, two events are triggered: (i) the membrane potential is reset to *V_r_* = 10 mV and *V_j_* is held fixed for a refractory period *τ*_r_ = 0.5 ms; (ii) a spike is emitted and received *τ_d_* = 0.55 ms later by the post-synaptic cells connected to neuron *j*. All the other parameters are initially set as in Ref. [23], namely: *b* = 0.8, *K* = 1000, *g* = 5, and *N* = 10,000.

We first compute the instantaneous probability density *P*(*v*) of membrane potentials *V_i_* ∈ [*v*, *v* + *dν*] for *J* = 0.1 mV and 0.5 mV. The asynchronous regime is by definition characterized by a constant firing rate [29] (in the thermodynamic limit). This implies that the flux of neurons along the *v*-axis is independent of both potential and time, i.e. the corresponding probability density *P*(*v*) is stationary. From Fig. 1, where three different snapshots of *P*(*v*) are plotted, we notice instead strong fluctuations, which appear to grow with the coupling strength *J*.

**FIG. 1.**
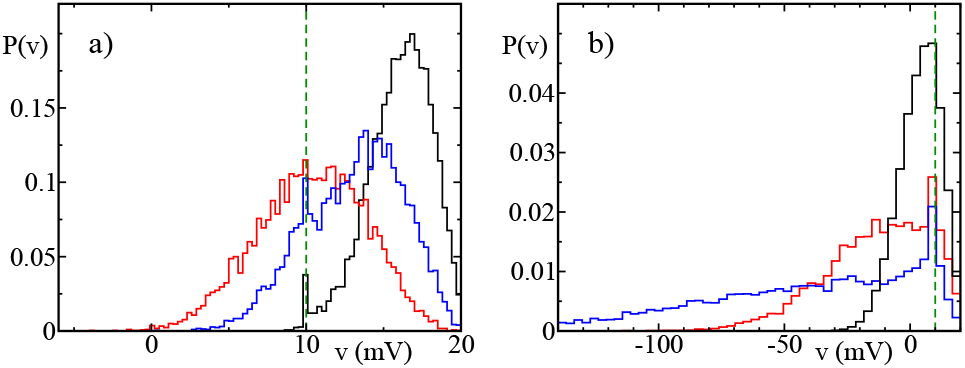
Three different instantaneous probability distributions of the membrane potentials *P*(*v*) for *N* = 10,000 and *J* = 0.1 mV (a) and *J* = 0.5 mV (b). The green dashed line refers to the reset potential *V*_r_.

Such large fluctuations are inconsistent with the stationarity of the asynchronous regime. In order to better understand their nature, it is necessary to take the thermodynamic limit. This can be done in various ways. In Ref. [21], *N* is let diverge to ∞, keeping all other parameters constant. This limit is not able to capture the fluctuations seen in Fig. 1, which indeed slowly vanish upon increasing *N*. In most of the literature on balanced states [10, 12-14], first the limit *N* → ∞, is taken, and then the average in-degree *K* is let diverge under the assumption that the coupling strength *J* is on the order of 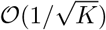, i.e. one can rewrite explicitly 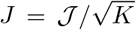, and 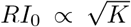. In this Article we propose to let *N* and *K* diverge simultaneously, assuming *K* = *cN* (this corresponds to assuming a massive connectivity). A priori, there are two meaningful setups that can be considered: (W) weak external current, which corresponds to assume that *RI*_0_ is independent of *N* (and thereby *K*); (S) strong external current, i.e. 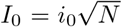. In the (W) setup the balance must be ensured a priori by imposing that excitatory and inhibitory fields nearly compensate each other. This is obtained by setting 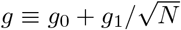 with *g*_0_ = *b*/(1 − *b*) so that the average difference between the excitation and inhibition is of the same order as statistical fluctuation. In the (S) setup there is no need to tune *g* because the external current *RI*_0_ maintains the balance. In this Article we show that CID emerges in both setups.

We first report the results for increasing network sizes for the setup (W), starting from *N* = 10,000, and including 40,000, 160,000, and 640,000. We set *c* = 0.1 and *g*_1_ = 100, as the resulting model, for *N* = 10,000, is equivalent to that in Ref. [23]. In Fig. 2a we plot the value of the average firing rate 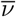 versus 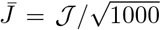 [30]. In order to damp the (small) sample-to-sample fluctuations, the results are averaged over seven, three and two different realizations of the network for *N* = 10, 000, 40, 000 and 160, 000, respectively. We observe a slow but clear convergence to an asymptotic curve in the entire range of coupling values. Finite-size corrections are negligible for 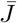 up to 0.1 mV, while for stronger coupling, the larger the network, the stronger is the tendency of the firing rate to decrease with the system size. Nevertheless, for *N* ≳ 160, 000 an asymptotic curve is attained, which exhibits a growth of 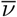 with 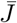 for sufficiently large coupling (compare with the solid full line, obtained by invoking the theoretical formula for a noise-driven LIF [31]). Another aspect that is maintained in the thermodynamic limit is a bursting activity, characterized by a coefficient of variation larger than 1 [32], for 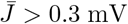 (data not shown).

**FIG. 2.**
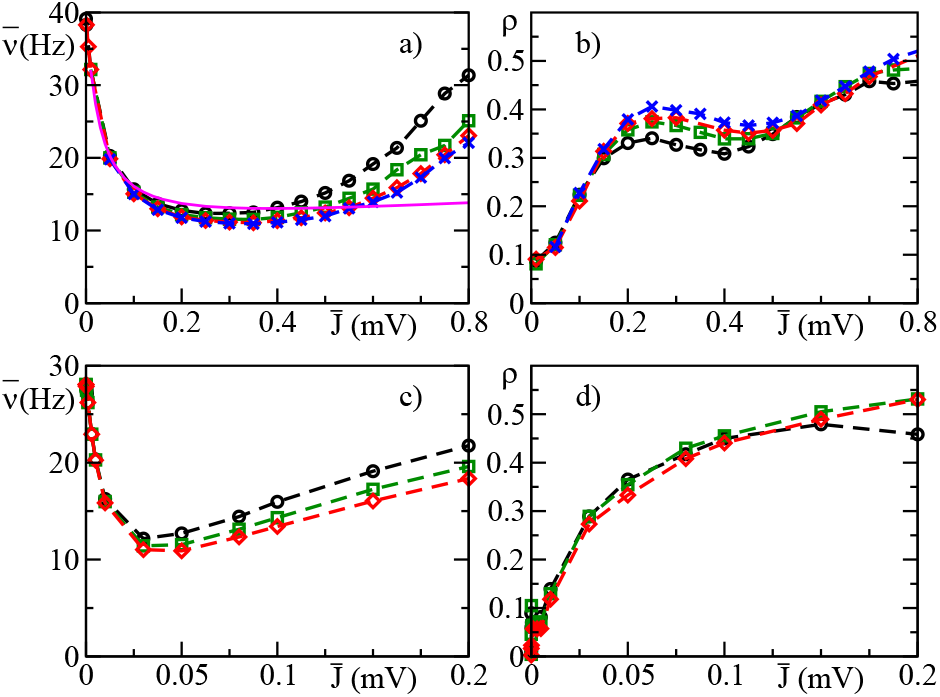
Average firing rate 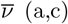 and order parameter *ρ* (b,d) versus the coupling strength 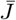 for different network sizes *N*: black circles refer to *N* = 10,000, green squares to 40,000, red diamonds to 160, 000 and blue crosses to 640, 000 (only in (a) and (b)). The results refer to the (W) setup for LIF (a,b) and QIF (c,d). The indicators have been averaged over a time window of 40 s and over up to seven realizations of the network. The solid magenta line (a) is obtained by invoking the diffusion approximation for a noise driven LIF [31].

A typical order parameter that is used to quantify the strength of collective dynamics is based on the relative amplitude of the macroscopic fluctuations [26]

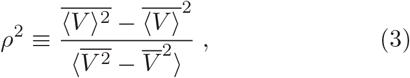

where 〈·〉 denotes an ensemble average, while the overbar is a time average. In practice *ρ* is the rescaled amplitude of the standard deviation of the average (*V*). When all neurons behave in exactly the same way (perfect synchronization), the numerator and the denominator are equal to one another and *ρ* = 1. If instead, they are independent, 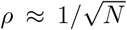. From the results plotted in Fig. 2b, we see that the order parameter *ρ* is finite in the whole range of the considered coupling. Furthermore it is substantially independent of *N* for 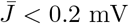, while for larger 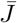 it exhibits a slower convergence to values ≃ 0.4 − 0.5. This clearly indicates that the thermodynamic phase is not a standard asynchronous regime, but is rather characterized by a collective dynamics, also for very small coupling strengths.

The nature of the macroscopic dynamics can be appreciated from the spectrum 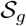 of the global activity *Y*(*t*) (obtained by summing the signals emitted by all the neurons) – for different system sizes. In Fig. 3 we plot the rescaled spectrum 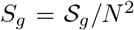 (panels a and b refer to 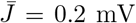 and 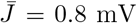, respectively). The data collapse suggests that the dynamics remains irregular in the thermodynamic limit, i.e. that the fluctuations are not finite-size effects. In fact, an asynchronous regime would have been characterized by a spectral amplitude 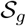 of order 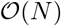) rather than 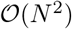). For both coupling strengths the spectral density is mostly concentrated in two frequency ranges: (i) around f ≈ 1800 Hz, which corresponds to the inverse of the delay; (ii) at low frequencies in a range that approximately corresponds to the firing rate. A relative comparison confirms that the collective dynamics is stronger for larger coupling strengths.

**FIG. 3.**
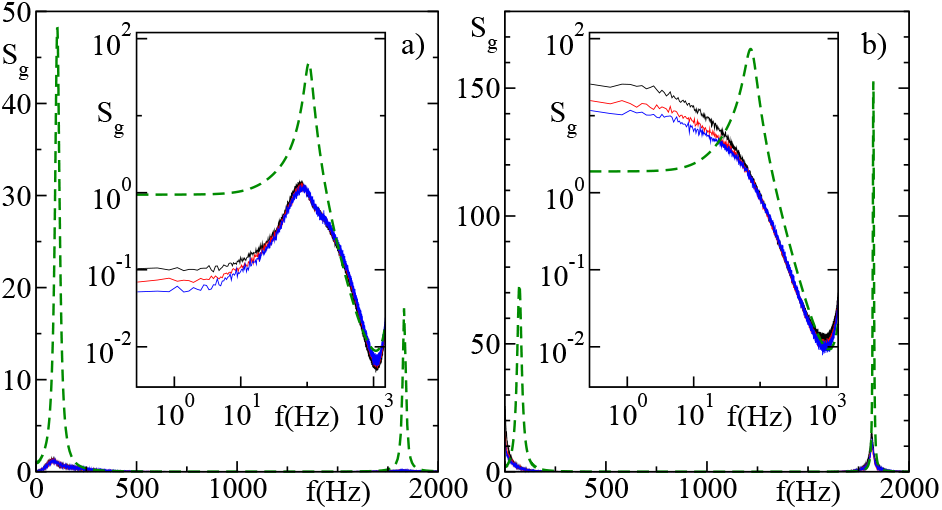
Global spike-train spectra *S_g_* (*f*) versus the frequency for 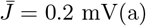 and 0.8 mV(b) for LIF in (W) setup. The different lines refer to different system sizes, namely *N* = 10,000 (black), *N* = 40, 000 (red) and *N* = 160, 000 (blue). The dashed green lines show the theoretical results obtained by following Ref. [21].

Altogether the broad-band structure of the spectrum suggests that the nature of the CID is stochastic-like even though the model is perfectly deterministic. The highdimensional character of the neural activity is confirmed by a fractal-dimension analysis [33, 34]. To our knowledge we provide the first convincing evidence of collective irregular behaviour in a balanced state. The closest regime is reported in a preprint [35], which deals with a fully coupled rate model.

The persistence of collective dynamics in the thermodynamic limit in the (W) setup can be understood in the limit of small connectivity, by revisiting the theory developed by Brunel [21] in the context of highly-diluted networks, under the implicit assumption that the thermodynamic limit is taken by letting the number of neurons diverge. The central point is the evolution equation for the probability *P*(*v*, *t*)

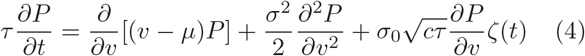

This stochastic Fokker-Planck equation was derived in [21] (see Eq. (32), here rewritten in our notations); it is valid so long as the current *I* can be represented as the sum of a deterministic contribution *μ* and a noise of amplitude *σ*. Self-consistent formulas for *μ* and *σ* can be derived upon assuming an uncorrelated Poisson activity of the various neurons, obtaining *μ*(*t*) = *RI*_0_ + *K Jτ*(*b* − (1 − *b*)*g*)*ν*(*t* − *τ_d_*), and 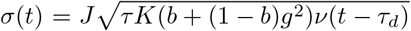 (see Eqs. (4,5) in [21]), where *ν*(*t*) is the instantaneous firing rate at time *t* [36].

The presence of the common noise in Eq. (4) is due to the sharing of a fraction of afferent neurons. In the context of [21], the additive noise is a finite-size effect, since *c* = *K*/*N* vanishes for *N* → ∞, while in our setup it remains finite. More precisely, *ζ*(*t*) is a white noise term - 〈*ζ*(*t*)*ζ*(*t* + *T*)〉 = δ(*T*) - while *σ*_0_ is the value of *σ* corresponding to the constant firing rate *ν*_0_ obtained within the diffusion approximation [31]. By inserting our scaling assumptions for *K*, *J*, and *g*, we find that in the thermodynamic limit 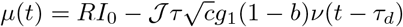 and 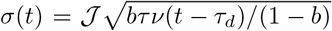, i.e. both parameters remain finite when the limit *N* → ∞ is taken. As a result, it makes sense to use the solution of Eq. (4) as a reference for the results of our numerical simulations. An analytic expression for the rescaled power spectrum can be found in [21] (see the expression reported at the end of page 204). The resulting shape for our parameter values is reported in Fig. 3 (see the green dashed line in panels a and b). A qualitative agreement is observed for both 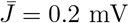 and 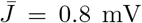, starting from the presence of a peak in correspondence of the inverse delay. The similarity between the low-frequency peaks is less pronounced for the higher coupling strength, showing that the true dynamics is definitely less regular than theoretically predicted by the noisy Fokker-Planck equation. The lack of a quantitative agreement is not a surprise, given the perturbative character of the noisy term and the assumption of a Poisson statistics that is not generally valid.

A closer agreement would be obtained if we could relax some of such approximations. A promising approach is the self-consistent method developed by Lindner and co-authors [37, 38], which provides a more accurate description of the spiking activity of LIF neurons. Unfortunately, for our setup above 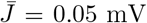 the method does not converge [39], thus leaving open the question whether it is a technical or conceptual matter.

We now discuss the strong-current setup (S), with 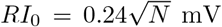 (i.e. *Ri*_0_ = 0.24 mV) and *g* = 5. In this case, the balance is attained (at leading order in *N*) by imposing the condition *I_i_* + *I*_0_ = 0. Under the assumption of a constant firing rate, this implies 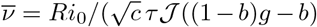. The results of simulations for 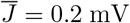 and different values of *N* are reported in Fig. 4, where one can see that the firing rate converges towards the expected asymptotic value 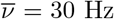, with a 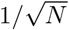 rate (see the solid line). More important is that the order parameter *ρ* remains finite for increasing *N* (see the inset). The presence of strong finite-size corrections prevent us from determining its asymptotic value; it is however clear that *ρ* does not vanish, indicating that a collective dynamics emerges also in the presence of strong external currents.

**FIG. 4.**
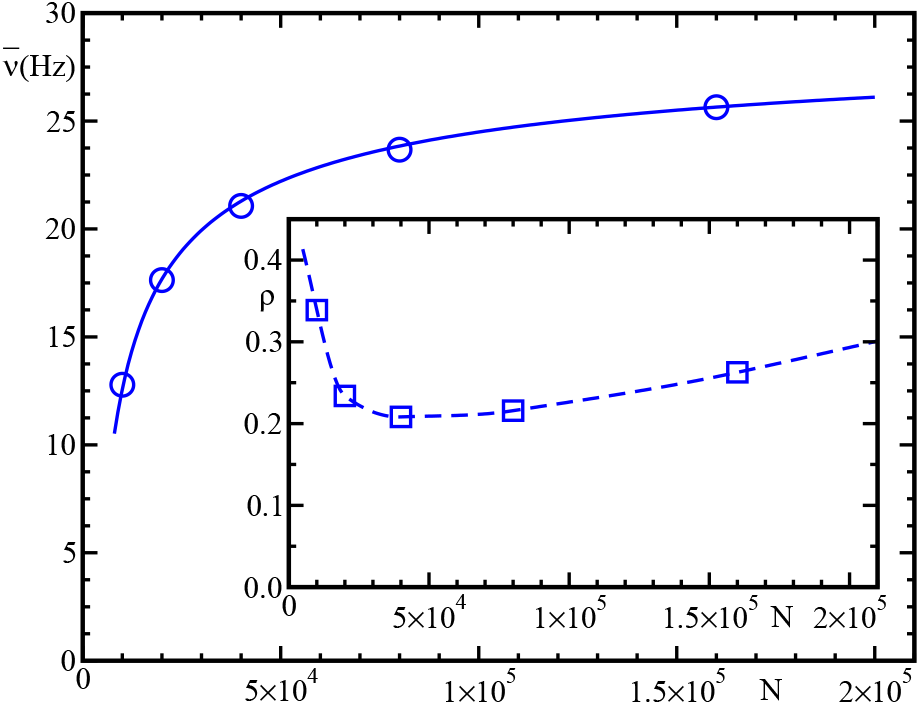
Mean firing rate 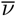 as a function of the system size *N* for 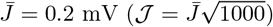 for the LIF in (S) setup. The solid line, 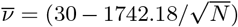, shows the convergence to the expected asymptotic value 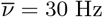. The inset shows the dependence of the synchronisation measure *ρ* on *N* (the dashed line is a guide for the eyes).

In order to establish the generality of CID in balanced, massively coupled networks, we have analysed another model, the QIF, which represents the canonical model for class I excitability [16, 40, 41]. Its evolution equation reads as

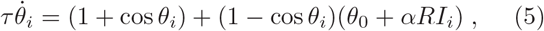

where *θ_i_* is an adimensional phase-like variable, *θ_th_* = *π* and *θ_r_* = −*π* are the threshold and reset value, respectively. Moreover, *θ*_0_ = 0.2, *α* = 1mV^−1^, while *RI_i_* is still defined as in Eq. (2), and all the other parameters are as for the LIF. As shown in Fig. 2c,d, where the firing rate 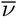 and the order parameter *ρ* are reported for different coupling strengths, there is again a clear evidence of synchronization. The broadband structure of the corresponding spectra of the neural activity (data not shown) indicate that the collective dynamics is stochastic-like.

Finally, we made several other tests, eliminating refractoriness, setting the delay equal to zero and adding noise to the external current (as in the original Brunel paper [21]). In all these cases *ρ* remains finite and exhibits an irregular behavior.

Altogether, we have found that CID emerges in all massively coupled networks we have explored. This comes as a surprise: in other models of massively coupled neuronal systems, the microscopic chaotic dynamics which may emerge in finite systems, disappears in the thermodynamic limit [42, 43], while here it does not only survive but contributes to sustain a macroscopic stochastic-like evolution. This point definitely needs to be better clarified.

The evidence that CID survives in the vanishing coupling limit could represent the starting point for future progress. In fact, for *J* = 0 any distribution *P*(*v*) is a marginally stable solution and can in principle be (de)stabilized by an arbitrarily small coupling. Such a singular behavior was successfully handled to explain the onset of partial synchrony [4], by mapping the ensemble of LIF neurons onto the much simpler Kuramoto-Daido equation [44, 45]. Can one hope to make a similar analysis in the context of the balanced regime?

The authors acknowledge N. Brunel, F. Farkhooi, G. Mato, M. Timme, and M. di Volo for enlightening discussions. This work has been partially supported by the French government under the A⋆MIDEX grant ANR-11-IDEX-0001-02 (A.T.) and mainly realized at the Max Planck Institute for the Physics of Complex Systems (Dresden, Germany) within the activity of the Advanced Study Group 2016/17 “From Microscopic to Collective Dynamics in Neural Circuits”.

